# Effects of exclusive breastfeeding on infant gut microbiota: a meta-analysis across studies and populations

**DOI:** 10.1101/292755

**Authors:** Nhan T. Ho, Fan Li, Kathleen A. Lee-Sarwar, Hein M. Tun, Bryan Brown, Pia S. Pannaraj, Jeffrey M. Bender, Meghan B. Azad, Amanda L. Thompson, Scott T. Weiss, M. Andrea Azcarate-Peril, Augusto A. Litonjua, Anita L. Kozyrskyj, Heather B. Jaspan, Grace M. Aldrovandi, Louise Kuhn

## Abstract

Literature regarding the differences in gut microbiota between exclusively breastfed (EBF) and non-EBF infants is meager with large variation in methods and results. We performed a meta-analysis of seven studies (a total of 1825 stool samples from 684 infants) to investigate effects of EBF compared to non-EBF on infant gut microbiota across different populations. In the first 6 months of life, overall bacterial diversity, gut microbiota age, relative abundances of Bacteroidetes and Firmicutes and microbial-predicted pathways related to carbohydrate metabolism were consistently increased; while relative abundances of pathways related to lipid, vitamin metabolism and detoxification were decreased in non-EBF vs. EBF infants. The perturbation in microbial-predicted pathways associated with non-EBF was larger in infants delivered by C-section than delivered vaginally. Longer duration of EBF mitigated diarrhea-associated gut microbiota dysbiosis and the effects of EBF persisted after 6 months of age. These consistent findings across vastly different populations suggest that one of the mechanisms of short and long-term benefits of EBF may be alteration in gut microbes.

## Background

Establishment of the gut microbiota in early life has substantial impact on subsequent health ^1^. Common sources of the infant’s intestinal microorganisms are from the mother’s skin, vagina, stool and from breastfeeding ^2–5^. There is a close relationship between the infant’s gut microbiota and the mother’s breast milk microbiota and human milk oligosaccharides (HMO) composition ^6–9^. Indeed, recent evidence has shown that breast milk microbiota can directly seed the infant gut microbiota and the effects of breastmilk on infant gut microbiota is dose-dependent ^5^. The microbiota in breast milk changes over time during lactation and has been shown to be different between exclusive breastfeeding (EBF) vs. non-EBF mothers ^10,11^. Gut microbial abundances in breast-fed infants, especially bifidobacterial species, are correlated with the mother’s HMOs and HMO-related catabolic activity ^3,12,13^. Infant gut microbiota have been shown to be different between breast fed vs. formula-fed infants ^14–20^ and change rapidly during the transition from breastfeeding to formula ^21^.

EBF in the first 6 months of life provides a multitude of health benefits ^22,23^. For example, EBF has been shown to be strongly protective against diarrhea morbidity and mortality ^24^ and decreases long-term risk of diabetes and obesity as compared to non-EBF or formula-fed infants ^25–27^. We hypothesize that the numerous benefits of EBF may be in part due to its effects on the infant gut microbiota. Several recent studies have identified varying differences in gut microbial composition or diversity between EBF and non-EBF infants ^5,28–30^ or gradients in the gut microbiota composition or diversity across EBF, non-EBF and non-breastfed (non-BF) infants ^5,14,29,31,32^. However, some other studies have found no significant differences in gut microbial communities between EBF and non-EBF infants ^3,33^. In addition, mode of delivery has been variably reported to have no effect ^34^ or a significant effect ^30^ or a potential interaction effect with breastfeeding ^31^ on the infant gut microbiota. The wide variation in reported results together with heterogeneity in feeding category definitions, study designs, study populations, and especially in data processing and analysis methods make these findings difficult to synthesize and interpret.

In this study, we applied appropriate and robust statistical methods to analyze gut microbiome data and performed meta-analyses pooling estimates from seven microbiome studies (a total of 1825 stool samples from 684 infants) to investigate the effects of EBF compared to non-EBF on infant gut microbiota in the first 6 months of life across different populations. In addition, we evaluated effect modification by infants’ mode of delivery. We aimed to synthesize existing data and to examine consistency or inconsistency in findings across studies and populations. Where data were available, we also examined persisting effects of EBF occurring in the first 6 months on the gut microbiota of infants after 6 months of life as well as interaction effects between diarrhea and EBF on the infant gut microbiota.

## Results

### Gut microbial diversity is increased in non-EBF vs. EBF infants ≤ 6 months of age

In infants ≤ 6 months of age, across the seven included studies, non-EBF consistently associated with increased gut microbial alpha diversity (standardized Shannon index) compared to EBF adjusting for infant age at sample collection (pooled standardized diversity difference (DD)= 0.34 standard deviation (sd), 95% confidence interval (95% CI)= [0.20; 0.48], pooled p-value<0.0001) (Figure 1a, Figure 1b). In a subset of five studies that also contained non-BF infants (Bangladesh ^35^, Canada ^31^, USA (California-Florida (CA-FL)) ^5^, USA (California-Massachusetts-Missouri (CA-MA-MO)) ^18^, and USA (North Carolina (NC)) ^29^), gut microbiome diversity (standardized Shannon index) was significantly increased in infants with less breastfeeding after adjusting for age of infants at sample collection (pooled standardized DD=0.39 sd, 95% CI=[0.19; 0.58], pooled p-value=0.0001) (Figure 1c). Results were consistent utilizing three other commonly used alpha diversity indices (Phylogenetic diversity whole tree, Observed species, Chao1) (all pooled p-values <0.05) (Figure 1d, Figure 1e).

**Figure 1.**
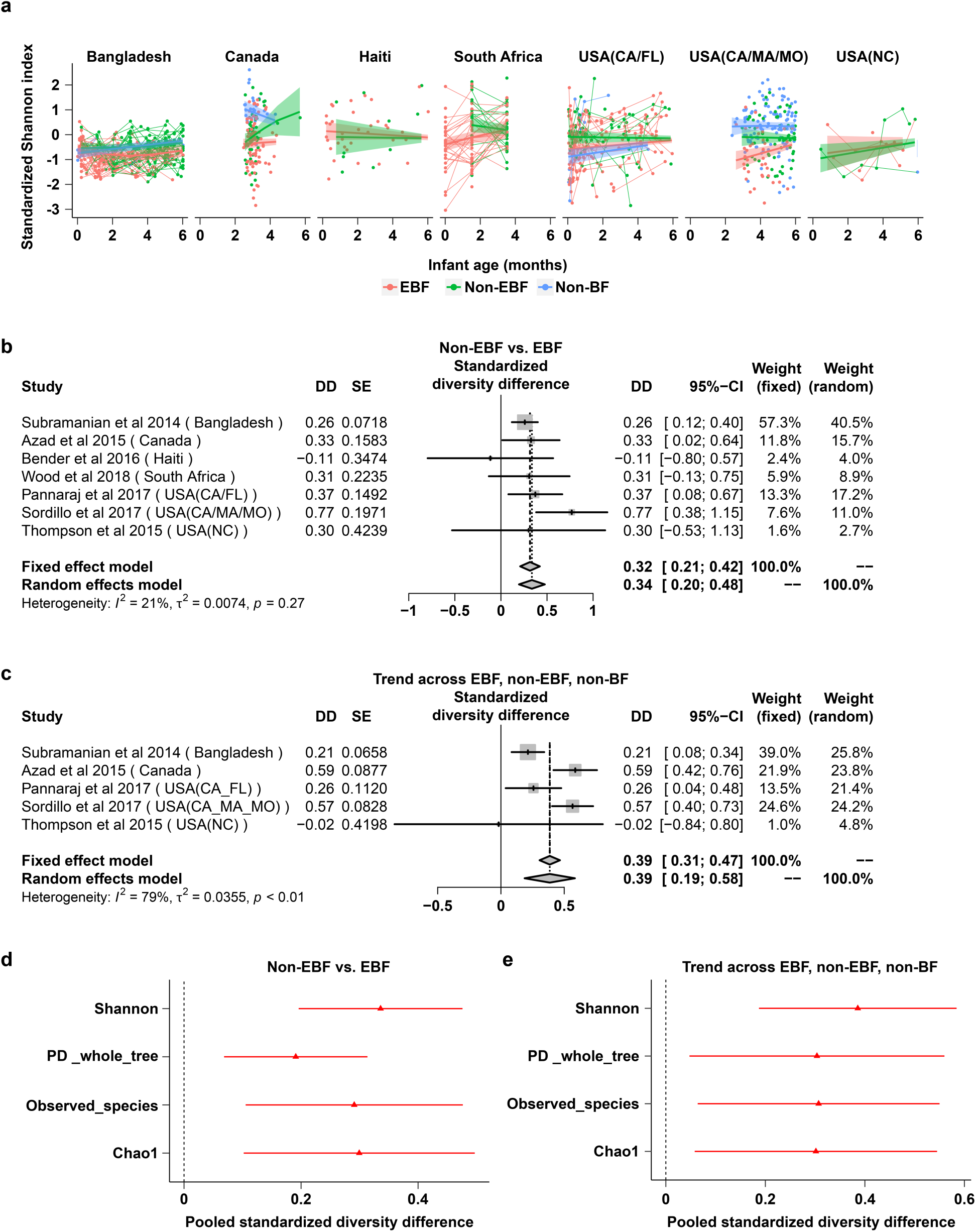
Meta-analysis: the effects of non-exclusive vs. exclusive breastfeeding on gut microbial diversity in infants ≤ 6 months of age. **a**: Gut microbial alpha diversity (standardized Shannon index) by breastfeeding status by age of infants at stool sample collection from each of seven included studies. Fitted lines and 95% confidence intervals (95% CI) were from Generalized Additive Mixed models (GAMM). **b**: The difference in gut microbial alpha diversity (standardized Shannon index) between non-exclusively breastfed (non-EBF) vs. exclusively breastfed (EBF) infants ≤ 6 months of age from each study and the pooled effect across seven included studies (meta-analysis). **c**: The trend effect of gut microbial alpha diversity (standardized Shannon index) across EBF, non-EBF and non-BF infants ≤ 6 months of age from each study and the pooled effect across five included studies (meta-analysis). Data from the Haiti and South Africa studies were not included as there was no non-BF group in these two studies. In each study, to roughly test for trends across breastfeeding categories, breastfeeding was coded as a continuous variable in the models (EBF=1, non-EBF=2 and non-BF=3). **d**: Pooled estimates and 95% CI for the difference in gut microbial alpha diversity (four common alpha diversity indexes (all standardized)) between non-EBF vs. EBF infants ≤ 6 months of age across seven studies. **e**: Pooled estimates and 95% CI for the trend effect of gut microbial alpha diversity (four common alpha diversity indexes (all standardized)) across EBF, non-EBF and non-BF infants ≤ 6 months of age. Estimates for diversity difference or trend and corresponding standard errors from each study were from linear mixed effect models (for longitudinal data) or linear models (for non-longitudinal data) and were adjusted for age of infants at sample collection. Pooled estimates of standardized diversity difference or trend and their 95% CI were from random effect meta-analysis models based on the adjusted estimates and corresponding standard errors of all included studies. Pooled estimates with pooled p-values<0.05 are in red and those with false discovery rate (FDR) adjusted pooled p-values <0.1 are in triangle shape. EBF: exclusive breastfeeding; non-EBF: non-exclusive breastfeeding; non-BF: non-breastfeeding; USA: United States of America; CA: California; FL: Florida; MA: Massachusetts; MO: Missouri; NC: North Carolina; DD: Diversity difference; SE: Standard error.

In sensitivity meta-analyses excluding estimates from either the USA (NC) study ^29^ (which contained a small number of infants ≤ 6 months old), or the Haiti study ^3^ (which included samples from HIV-uninfected infants born to HIV-infected and HIV-uninfected mothers) or the Vitamin D Antenatal Asthma Reduction Trial (VDAART) study ^18^ in the USA (CA-MA-MO) (which contained samples from infants at high risk of asthma and allergies, half of whom were randomized to high-dose antenatal vitamin D supplementation), the results remained similar (Supplementary figure 1).

In a subset of four studies (Canada ^31^, Haiti ^3^, USA (CA-FL) ^5^, and USA (CA-MA-MO) ^18^) with available data on mode of delivery, the increase in microbial diversity associated with non-EBF was similar in the meta-analysis stratified on vaginally-delivered infants and on cesarean-delivered infants (Supplementary figure 2).

**Figure 2.**
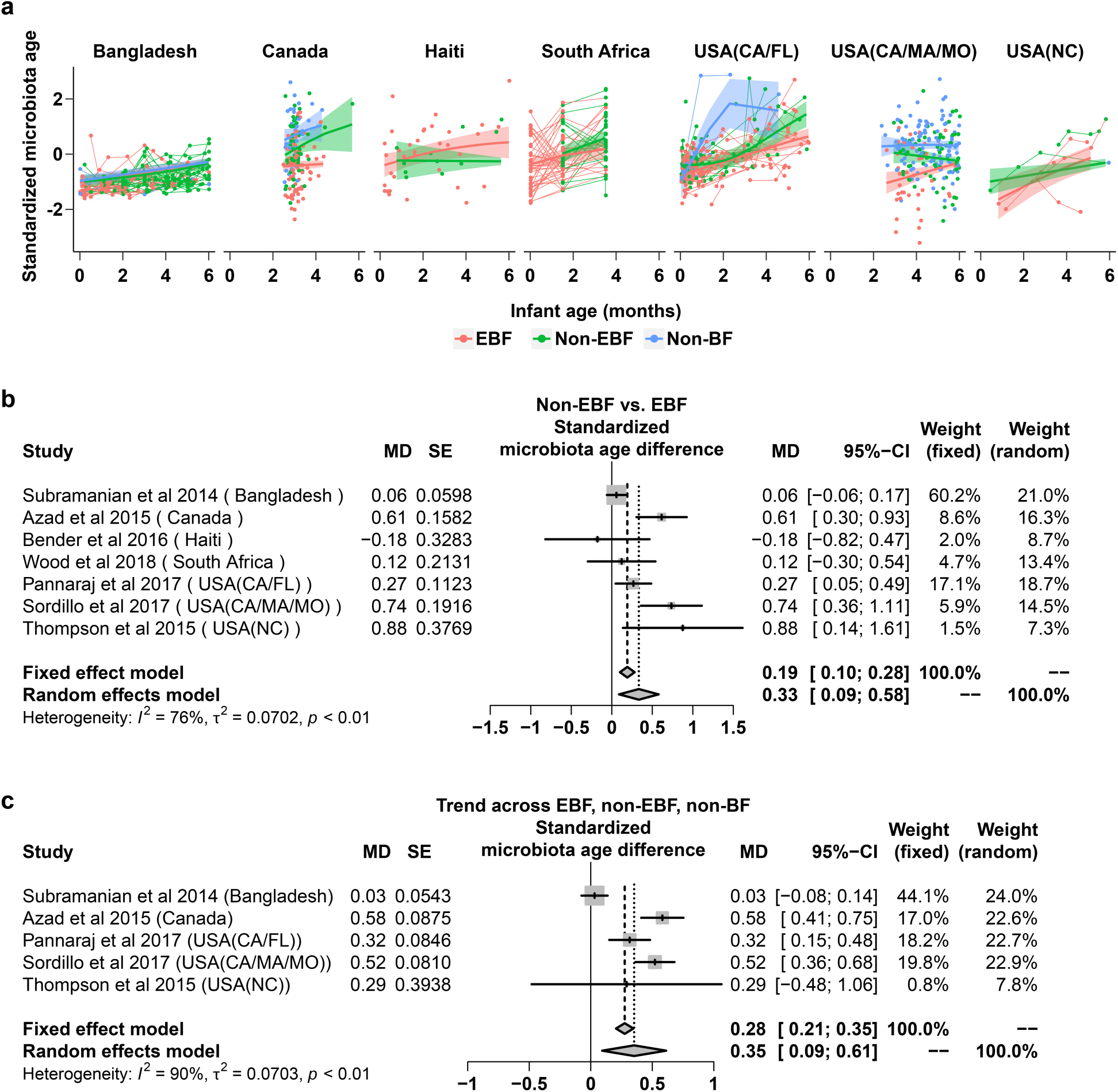
Meta-analysis: the effects of non-exclusive vs. exclusive breastfeeding on gut microbiota age in infants ≤ 6 months of age. **a**: Gut (standardized) microbiota age of infants ≤ 6 months of age by breastfeeding status by age of infants at stool sample collection from each of seven included studies. Fitted lines and 95% confidence intervals (95% CI) were from Generalized Additive Mixed models (GAMM). **b**: The difference in gut (standardized) microbiota age between non-exclusively breastfed (non-EBF) vs. EBF infants ≤ 6 months of age from each study and the pooled effect across seven included studies (meta-analysis). **c**: The trend of gut (standardized) microbiota age across EBF, non-EBF and non-BF infants ≤ 6 months of age from each study and the pooled effect across five included studies (meta-analysis). The Haiti and South Africa studies were not included as there was no non-BF group in these two studies. In each study, to test for trend across breastfeeding categories, breastfeeding was coded as a continuous variable in the model (EBF=1, non-EBF=2 and non-BF=3). Estimates for (standardized) microbiota age difference or trend and corresponding standard error from each study were from linear mixed effect models (for longitudinal data) or linear models (for non-longitudinal data) and were adjusted for age of infants at sample collection. EBF: exclusive breastfeeding; non-EBF: non-exclusive breastfeeding; non-BF: no breastfeeding; USA: United States of America; CA: California; FL: Florida; MA: Massachusetts; MO: Missouri; NC: North Carolina; MD: Microbiota age difference; SE: Standard error.

### Gut microbiota age is increased in non-EBF vs. EBF infants ≤ 6 months of age

A Random Forest (RF) model was used to predict the infant age in each included study based on relative abundances of the shared gut bacterial genera of the seven included study (Supplementary table 1). The model explained 95% of the variance related to chronologic age in the training set and 65% of the variance related to chronologic age in the test set of Bangladesh data (Supplementary figure 3). The predicted infant age in each included study based on relative abundances of the shared gut bacterial genera using this Random Forest model was regarded as gut microbiota age.

**Table 1.**
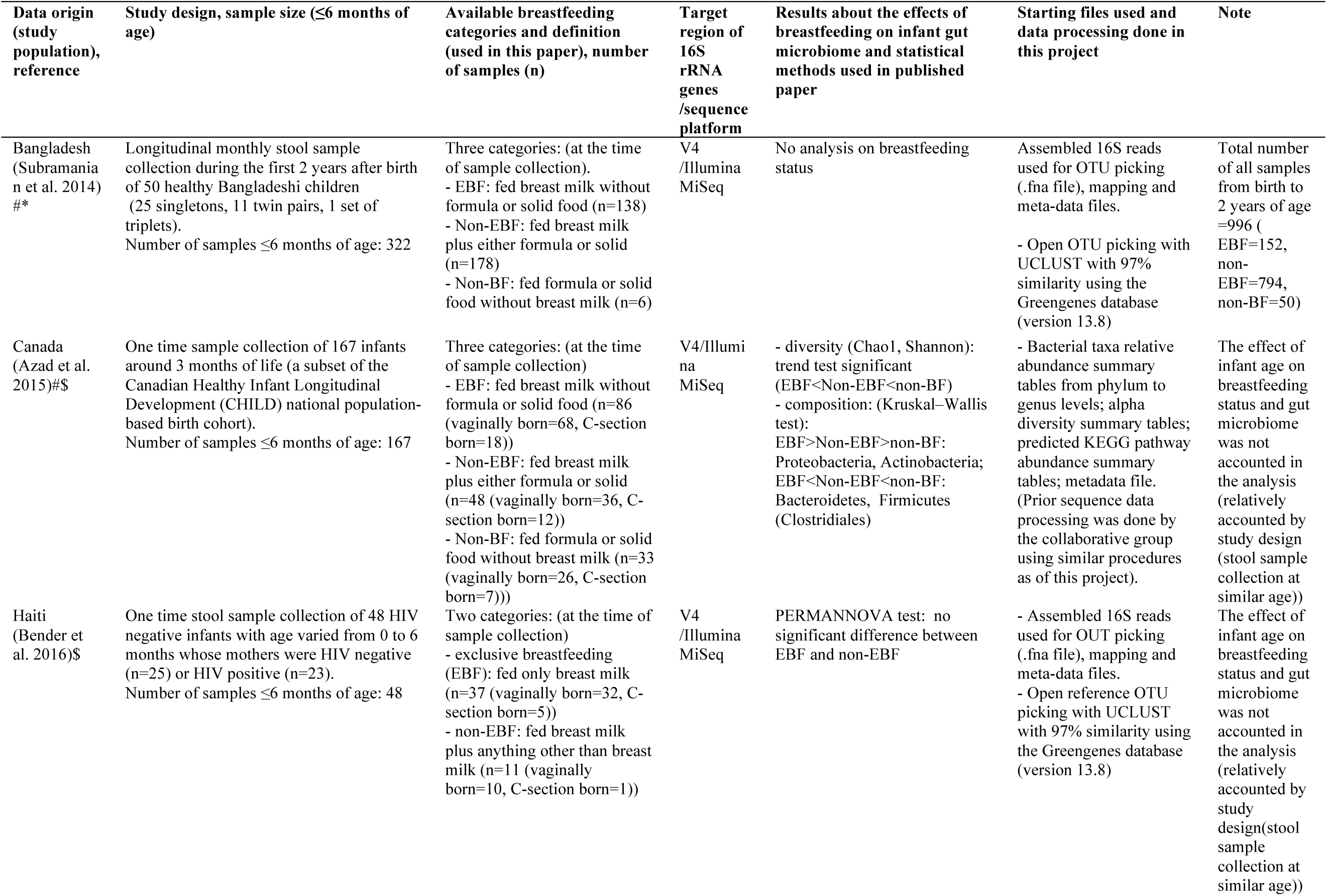
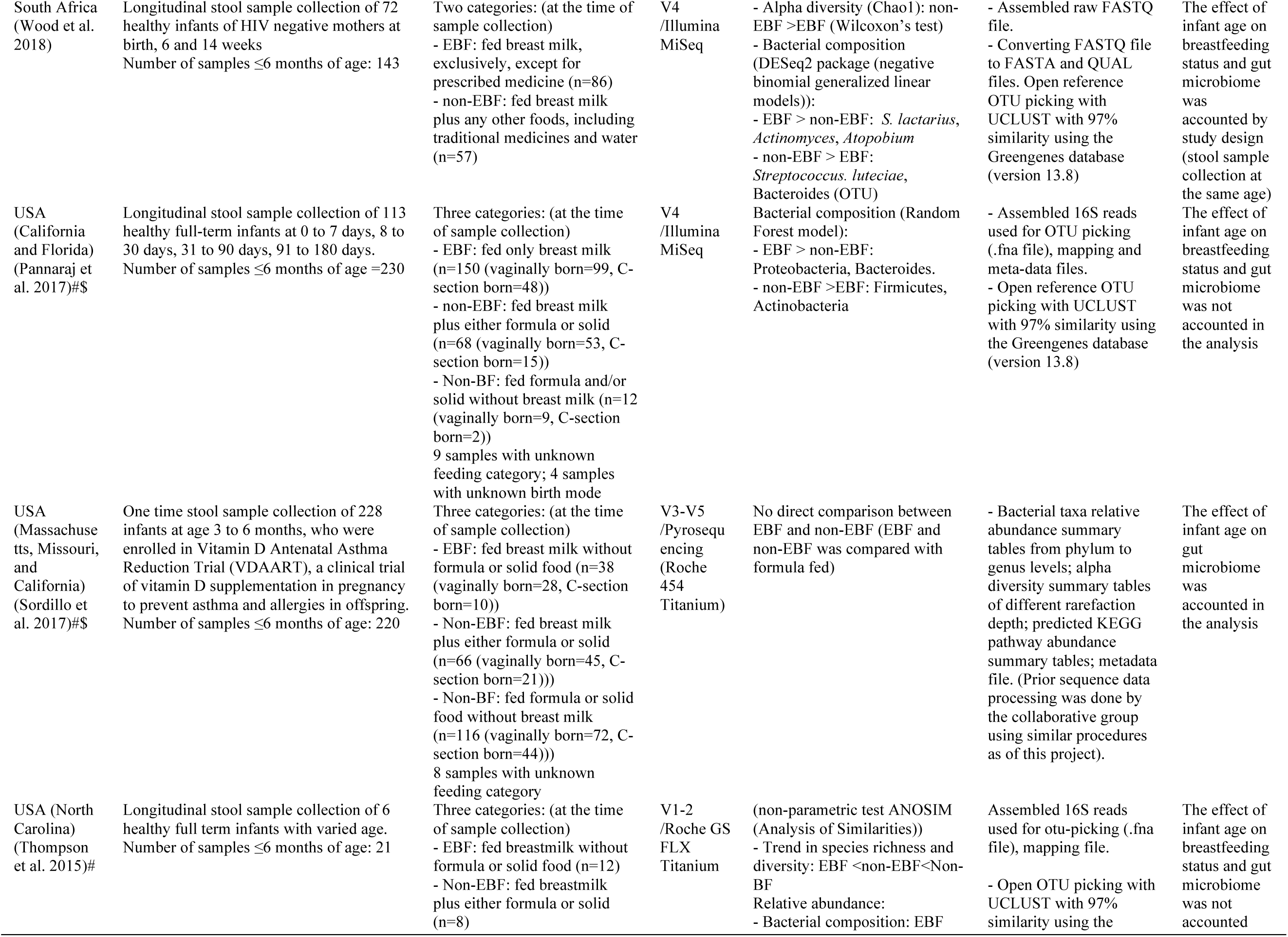
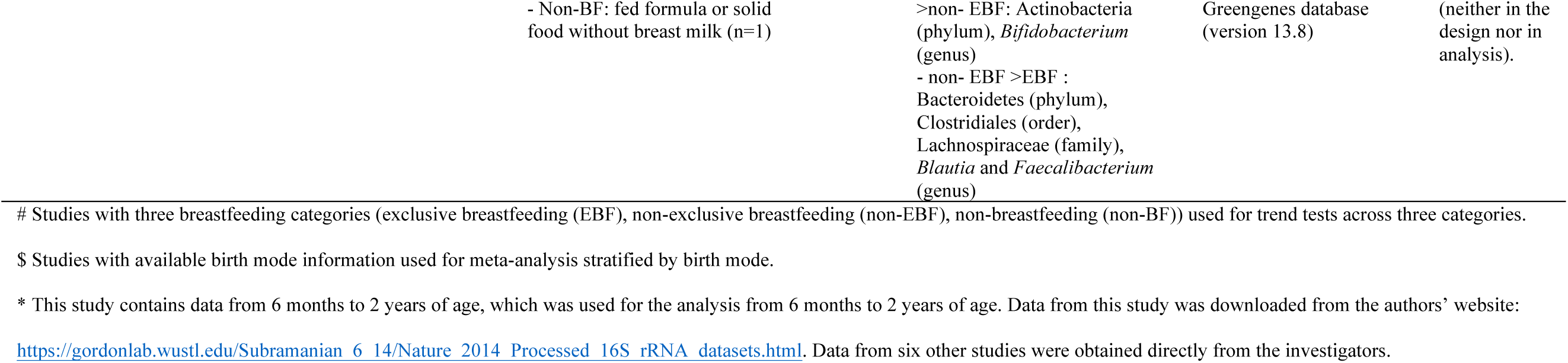
Summary of included studies.

**Figure 3.**
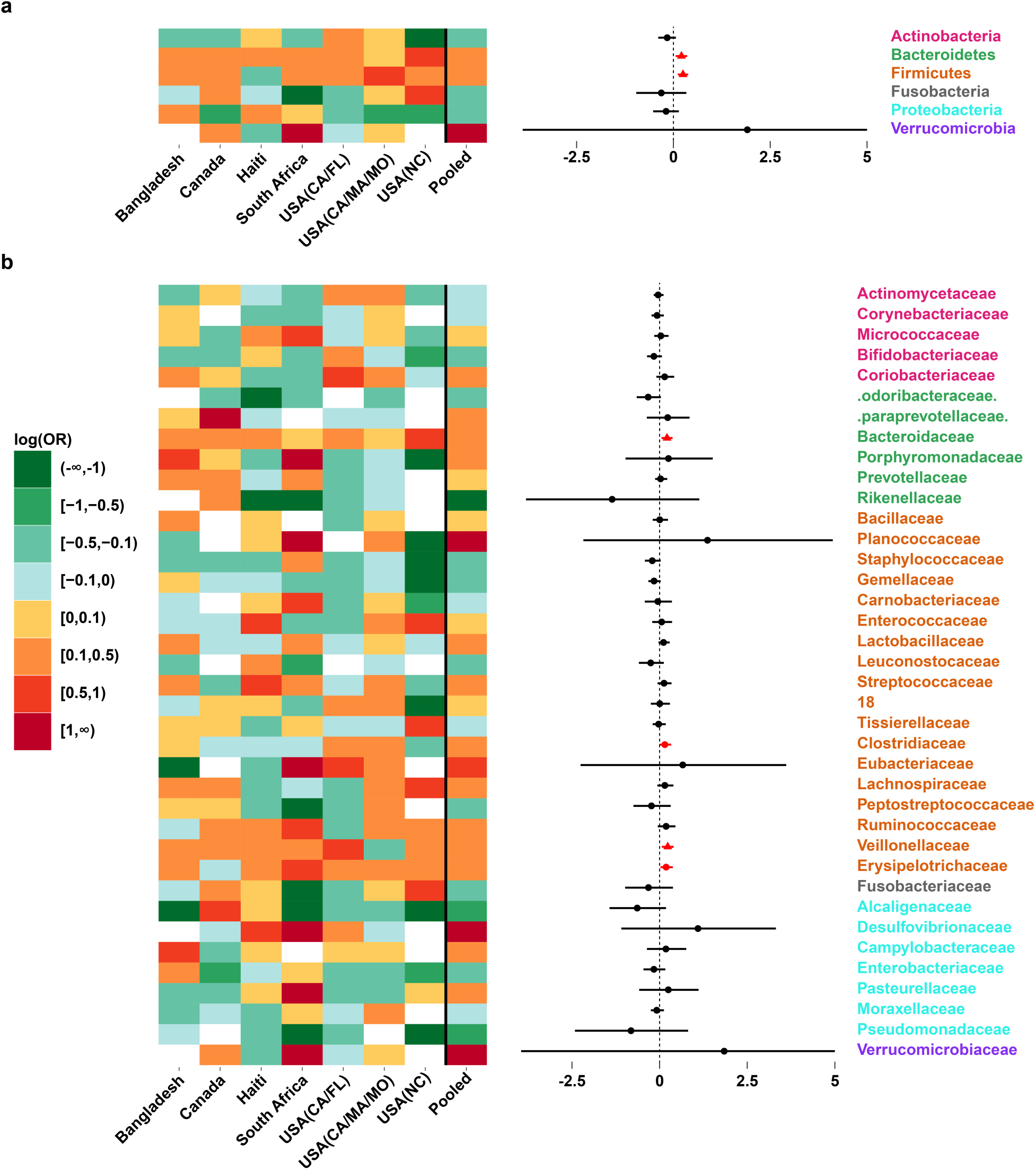
Meta-analysis: the effects of non-exclusive vs. exclusive breastfeeding on relative abundances of gut bacterial taxa in infants ≤ 6 months of age. **a**: Gut bacterial phyla: heatmap of log(odds ratio) (log(OR)) of relative abundances of all gut bacterial phyla between non-exclusively breastfed (non-EBF) vs. exclusively breastfed (EBF) infants for each study and pooled estimates across all seven studies with 95% confidence intervals (95% CI) (forest plot). **b**: Gut bacterial families: heatmap of log(OR) of relative abundances of all gut bacterial families between non-EBF vs. EBF infants for each study and pooled estimates across all seven studies with 95% CI (forest plot). All log(OR) estimates of each bacterial taxa from each study were from Generalized Additive Models for Location Scale and Shape (GAMLSS) with zero inflated beta family (BEZI) and were adjusted for age of infants at sample collection. Pooled log(OR) estimates and 95% CI (forest plot) were from random effect meta-analysis models based on the adjusted log(OR) estimates and corresponding standard errors of all included studies. Pooled log(OR) estimates with pooled p-values<0.05 are in red and those with false discovery rate (FDR) adjusted pooled p-values <0.1 are in triangle shape. Missing (unavailable) values are in white. Bacterial families with unassigned name are numbered. USA: United States of America; CA: California; FL: Florida; MA: Massachusetts; MO: Missouri; NC: North Carolina.

In infants ≤ 6 months of age, a consistent (6/7 studies) increase of gut microbiota age was observed in non-EBF as compared to EBF infants after adjusting for infant age at sample collection (pooled standardized microbiota age difference (MD)= 0.33 sd, 95%CI= [0.09; 0.58], pooled p-value= 0.0074) (Figure 2a, Figure 2b). In sensitivity meta-analyses excluding either of the three studies mentioned above ^3,18,29^, the association remained unchanged (Supplementary figure 4). In the subset of four studies with available data on mode of delivery ^3,5,18,31^, meta-analysis stratified on vaginally-delivered infants and on cesarean-delivered infants showed similar overall increase in microbiota age in non-EBF vs. EBF infants (Supplementary figure 5).

The trend of increasing gut microbiota age in infants with less breastfeeding after adjusting for age of infants at sample collection was also observed in a subset of five studies containing a non-BF group ^5,18,29,31,35^ (pooled standardized MD=!0.35 sd, 95%CI= [0.09; 0.61], pooled p-value =0.0079) (Figure 2c).

**Figure 4.**
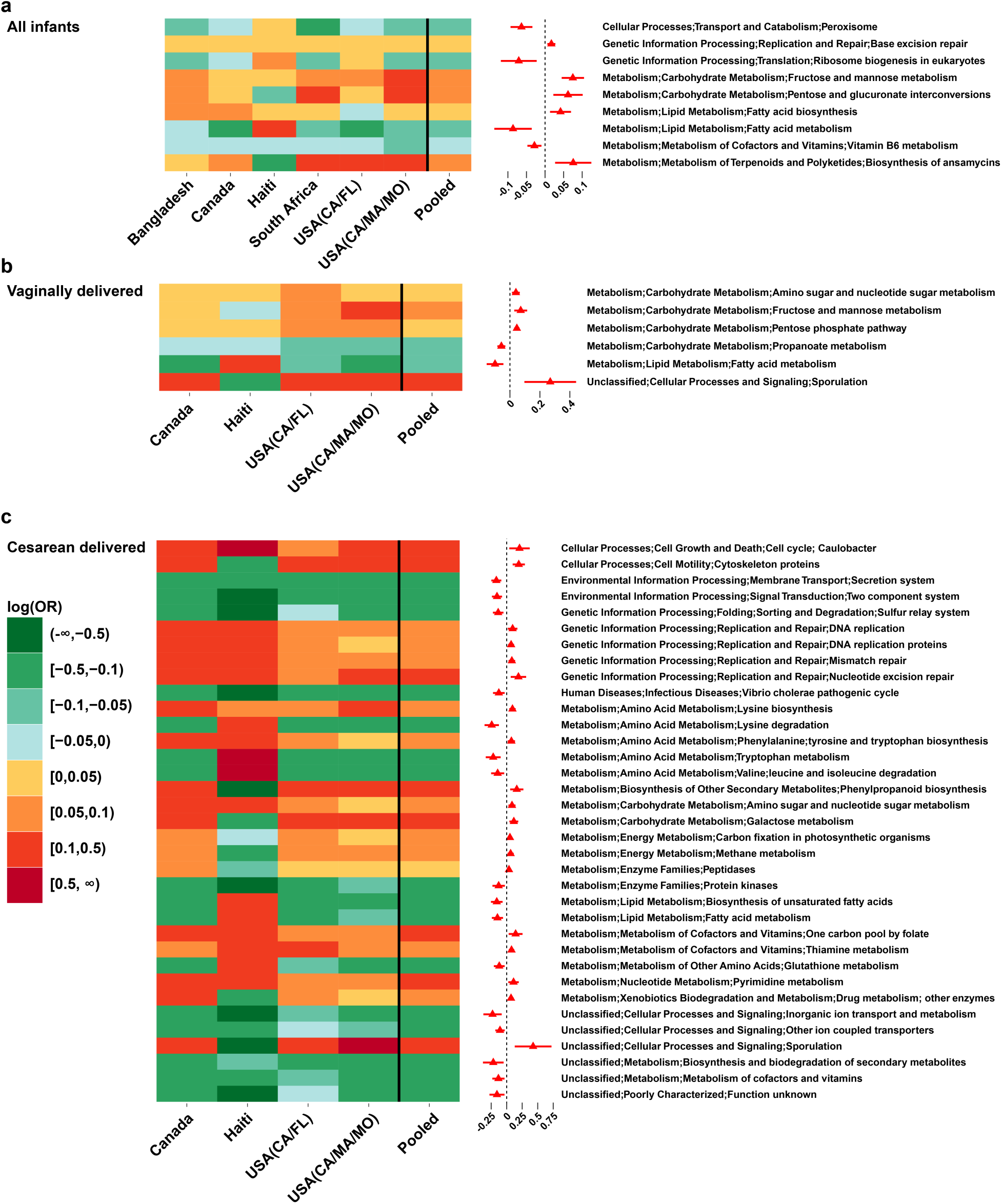
Meta-analysis: the effects of non-exclusive vs. exclusive breastfeeding on relative abundances of gut microbial KEGG (Kyoto Encyclopedia of Genes and Genomes) pathways in infants ≤ 6 months of age. **a**: Meta-analysis of all infants in all seven included studies: heatmap of log(odds ratio) (log(OR)) of relative abundances of gut microbial KEGG pathways at level 3 between non-exclusively breastfed (non-EBF) vs. exclusively breastfed (EBF) infants for each study and pooled estimates of all seven studies with 95% confidence intervals (forest plot). **b**: Meta-analysis of vaginally born infants in four studies: heatmap of log(OR) of relative abundances of gut microbial KEGG pathways at level 3 between non-EBF vs. EBF infants for each study and pooled estimates of four studies with 95% confidence intervals (forest plot). Only four studies with available birth mode information (Canada, Haiti, USA(CA_MA_MO) and USA(CA_FL)) are included. **c**: Meta-analysis of C-section born infants in four studies: heatmap of log(OR) of relative abundances of gut microbial KEGG pathways at level 3 between non-EBF vs. EBF infants for each study and pooled estimates of four studies with 95% confidence intervals (forest plot). Only four studies with available birth mode information (Canada, Haiti, USA(CA_MA_MO) and USA(CA_FL)) are included. All log(OR) estimates of each pathway from each study were from Generalized Additive Models for Location Scale and Shape (GAMLSS) with zero inflated beta family (BEZI) and were adjusted for age of infants at sample collection. Pooled log(OR) estimates and 95% confidence intervals (95%CI) (forest plot) were from random effect meta-analysis models based on the adjusted log(OR) estimates and corresponding standard errors of all included studies. Pooled log(OR) estimates with pooled p-values<0.05 are in red and those with false discovery rate (FDR) adjusted pooled p-values <0.1 are in triangle shape. Only pathways with FDR adjusted pooled p-value <0.1 are shown. USA: United States of America; CA: California; FL: Florida; MA: Massachusetts; MO: Missouri; NC: North Carolina.

**Figure 5.**
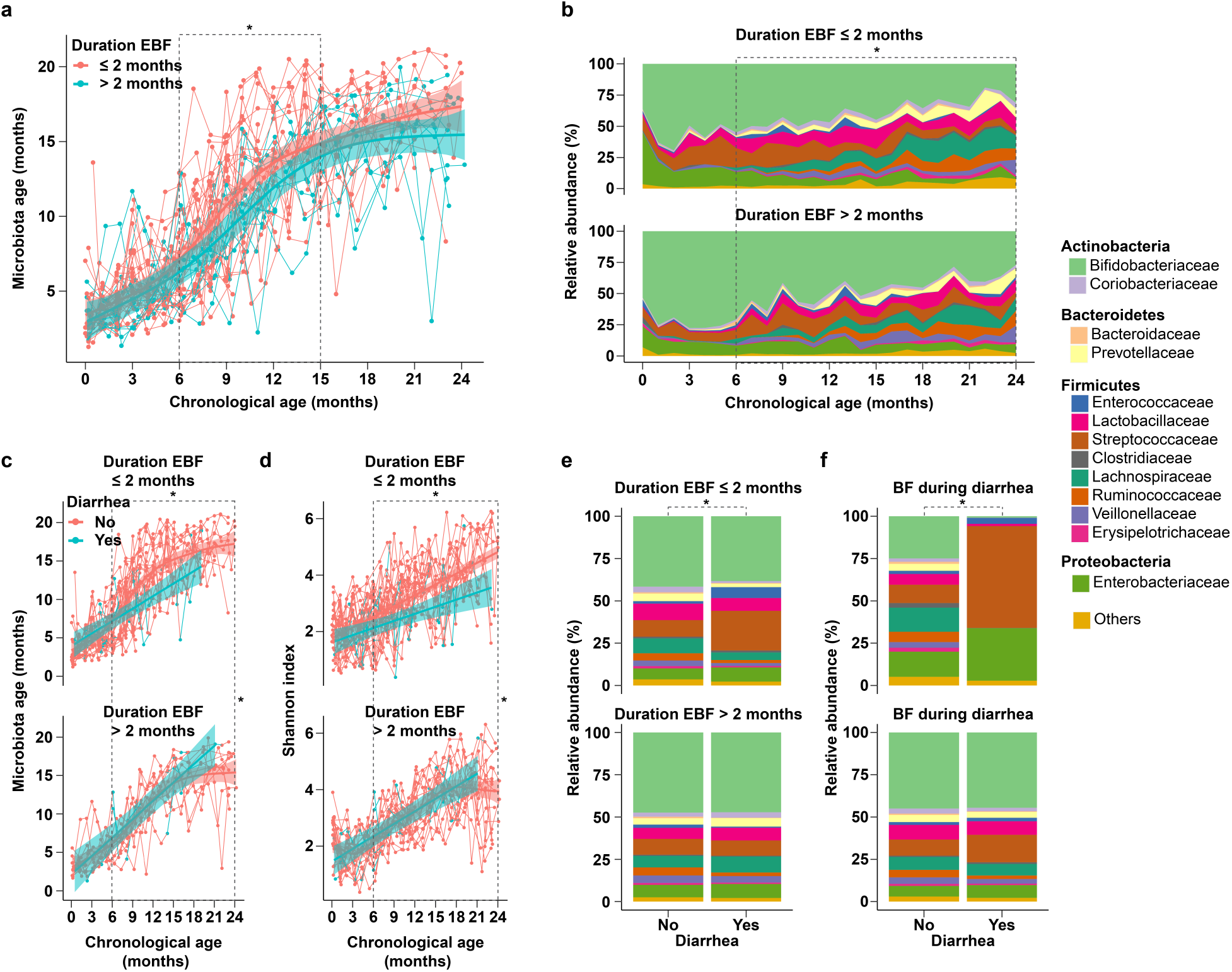
The continued effects of exclusive breastfeeding in the first 6 months of age on the infant gut microbiota up to 2 years of age. Data from Bangladesh study only. ***Upper panel***: The impact of duration of exclusive breastfeeding (EBF) on the gut microbiota of infants up to 2 years of age. **a**: the impact of duration of EBF shorter than 2 months vs. longer than 2 months from birth on infant gut microbiota age. **b**: the impact of duration of EBF shorter than 2 months vs. longer than 2 months from birth on infant gut bacterial family composition. Number of infants=50 (duration of EBF≤2 months=30, duration of EBF>2 months=20). Number of samples 0-2 years of age=996 (duration of EBF≤2 months=580, duration of EBF>2 months=416) ***Lower panel***: EBF mitigates diarrhea-associated gut microbiota dysbiosis in infants from 6 months to 2 years of age. **c**: the effects of diarrhea (vs. no diarrhea) around the time of stool sample collection on gut microbiota age in infants with duration of EBF shorter than 2 months vs. infants with duration of EBF longer than 2 months. **d**: the effects of diarrhea (vs. no diarrhea) around the time of stool sample collection on gut microbial diversity (Shannon index) in infants with duration of EBF shorter than 2 months vs. infants with duration of EBF longer than 2 months. **e**: the effects of diarrhea (vs. no diarrhea) on bacterial taxa composition at the family level around the time of stool sample collection in infants with duration of EBF shorter than 2 months vs. infants with duration of EBF longer than 2 months. **f**: the effects of diarrhea (vs. no diarrhea) on bacterial taxa composition at the family level around the time of stool sample collection in infants receiving no breastfeeding at the time of diarrhea vs. infants receiving breastfeeding at the time of diarrhea. Number of samples 6 months to 2 years of age= 674 (duration of EBF ≤2 months= 378 (diarrhea=29, no diarrhea=349); duration of EBF >2 months=296 (diarrhea=19, no diarrhea=277; with breastfeeding=616 (diarrhea=45, no diarrhea=571); without breastfeeding=44 (diarrhea=2, no diarrhea=42)). Fitted lines and 95% confidence intervals (95%CI) were from Generalized Additive Mixed models (GAMM). Dashed lines demarcate time periods tested. * indicate statistical significance.

### Gut microbial composition is altered in non-EBF vs. EBF infants ≤ 6 months of age

Across the seven included studies, there was a large heterogeneity in the difference (log(odds ratio)) of gut bacterial taxa relative abundances between non-EBF vs. EBF infants after adjusting for age of infants at sample collection. Notably, a decrease in relative abundance of Proteobacteria in non-EBF vs. EBF infants was observed in the four studies in North America but the opposite was observed in the other three studies in Haiti, South Africa and Bangladesh (Figure 3a). However, there was also remarkable consistency across studies. At the phylum level, there was an overall significant increase in the relative abundances of Bacteroidetes (consistent in all seven studies) and Firmicutes (consistent in 6/7 studies) in non-EBF vs. EBF infants (all pooled p-values <0.05 and false discovery rate (FDR) adjusted pooled p-values <0.1) (Figure 3a, Supplementary table 2). There was also a consistent trend of increasing relative abundances of these two phyla across EBF, non-EBF and non-BF in the subset of five studies with a non-BF group ^5,18,29,31,35^ (all pooled p-values <0.05 and FDR adjusted pooled p-values <0.1) (Supplementary table 3).

At the order level, the relative abundances of Bacteriodales (7/7 studies) and Clostridiales (6/7 studies) were consistently increased in non-EBF infants (all pooled p-values <0.05 and FDR adjusted pooled p-values <0.1) (Supplementary figure 6, Supplementary table 2). At the family level, the relative abundances of Bacteroidaceae (7/7 studies) and Veillonellaceae (6/7 studies) were consistently increased in non-EBF infants (all pooled p-values <0.05 and FDR adjusted pooled p-values <0.1) (Figure 3b, Supplementary table 2). At the genus level, there were increases in the relative abundances of *Bacteroides* (7/7 studies), *Eubacterium, Veillonella* (6/7 studies), and *Megasphaera* (5/7 studies) in non-EBF infants (all pooled p-values <0.05, FDR adjusted pooled p-value of *Eubacterium* <0.1) (Supplementary figure 7, Supplementary table 2). In sensitivity meta-analyses excluding either of the three studies mentioned above ^3,18,29^, results remained similar (Supplementary table 4, Supplementary table 5, Supplementary table 6).

In the subset of four studies with available data on mode of delivery ^3,5,18,31^, the results of meta-analysis stratified on vaginally-delivered infants were similar to those of meta-analysis stratified on cesarean-delivered infants from the phylum to family level. Phylum Proteobacteria (particularly family Enterobacteriaceae) was markedly and significantly reduced in non-EBF infants, especially among infants delivered by cesarean (Supplementary figure 8, Supplementary figure 9, Supplementary figure 10, Supplementary table 7). At the genus level, relative abundance of *Acidaminococcus* was significantly higher in vaginally-delivered non-EBF vs. vaginally-delivered EBF infants, whereas, relative abundances of *Proteus* and *Anaerotruncus* were significantly lower in cesarean-delivered non-EBF vs. cesarean-delivered EBF infants (all FDR adjusted pooled p-values <0.1) (Supplementary figure 11, Supplementary figure 12, Supplementary table 7).

### Gut microbial-predicted functions are perturbed in non-EBF vs. EBF infants ≤ 6 months of age

Across the seven included studies, although the difference (log(odds ratio)) of relative abundances of gut bacterial KEGG functional pathways between non-EBF vs. EBF infants was largely heterogeneous, important consistencies were found. At KEGG level two, there was no pathway significantly different between non-EBF and EBF group after adjusting for multiple testing (Supplementary figure 13, Supplementary table 8). At KEGG level three, the relative abundances of 24 pathways were significantly different between non-EBF and EBF infants (pooled p-values <0.05) and nine of these remained significant after adjusting for multiple testing (FDR adjusted pooled p-values <0.1) (Figure 4a). Specifically, after adjusting for age of infants at sample collection, in the non-EBF group vs. the EBF group, there were higher relative abundances of several pathways related to carbohydrate metabolism, viz. Fructose and Mannose Metabolism (7/7 studies), Pentose and Glucuronate Interconversions, and!Pentose Phosphate Pathway (6/7 studies), as well as Fatty Acid Biosynthesis and Biosynthesis of Ansamycins pathways (6/7 studies). In addition, in non-EBF infants, there were lower relative abundances of some pathways related to lipid metabolism, lipid homeostasis and free radical detoxification (Fatty Acid Metabolism (6/7 studies) and Peroxisome (6/7 studies)), Vitamin 6 Metabolism (7/7 studies) and Ribosome Biogenesis in Eukaryotes (5/7 studies) (all pooled p-values <0.05 and FDR adjusted pooled p-values <0.1) (Figure 4a, Supplementary table 8). Sensitivity meta-analyses excluding either of the three studies mentioned above ^3,18,29^ showed similar results (Supplementary table 9).

In the subset of four studies with available data on mode of delivery ^3,5,18,31^, meta-analysis stratified by mode of delivery showed remarkable heterogeneity between vaginally-delivered infants and cesarean-delivered infants. In vaginally-delivered infants, there were six pathways related to carbohydrate and lipid metabolism significantly perturbed between non-EBF vs. EBF infants after adjusting for multiple testing (all FDR adjusted pooled p-values <0.1) (Figure 4b, Supplementary table 10). Whereas, in cesarean-delivered infants only, there were a much larger number of pathways (35 pathways) of many cellular and metabolic processes (such as cell growth and death, membrane transport, replication and repair, carbohydrate, lipid, amino acid, vitamin, energy metabolism) significantly perturbed between non-EBF vs. EBF infants after adjusting for multiple testing (all FDR adjusted pooled p-values <0.1). In addition, the perturbation of these pathways were mostly consistent across four included studies (Figure 4c, Supplementary table 10).

### Duration of EBF in the first 6 months alters infant gut microbiota after 6 months of age

Using data from the Bangladesh study only, which included 996 stool samples collected monthly from birth to 2 years of life in 50 subjects, we found that shorter duration of EBF (less than 2 months vs. more than 2 months from birth) was associated with a larger increase in gut microbiota age. This association was significant from 6 months to 15 months of age (microbiota age difference (MD)= 1.64 months, 95%CI=(0.23, 3.05), p-value=0.0233) (Figure 5a).

From 6 months to 2 years of age, after adjusting for age of infants at sample collection, shorter duration of EBF (less than 2 months vs. more than 2 months from birth) was associated with lower relative abundance of the phylum Actinobacteria and higher relative abundance of Firmicutes (all FDR adjusted p-values <0.1) (Supplementary table 11). At the family level, infants with shorter duration of EBF had lower relative abundances of Bifidobacteriaceae, and Enterococcaceae and higher relative abundances of Lactobacillaceae, Coriobacteriaceae, Prevotellaceae, Clostridiaceae, Erysipelotrichaceae, and Lachnospiraceae (all FDR adjusted p-values <0.1) (Figure 5b, Supplementary table 11).

### Breastfeeding mitigates diarrhea-associated gut microbiota dysbiosis in infants from 6 months to 2 years of age

Again, using data from the Bangladesh study, we found that, after adjusting for age of infants at sample collection, diarrhea at the time of sample collection (vs. no diarrhea) was associated with reduced gut microbiota age in infants who received less than 2 months of EBF (MD= −1.17 months, 95% CI= (−2.11;-0.23), p= 0.0146) (Figure 5c). Diarrhea in infants who received less than 2 months of EBF was also associated with reduced gut microbial diversity as showed by Shannon index (diversity difference =-0.58, 95%CI=(−0.83,-0.34), p<0.0001) (Figure 5d) and the three other common alpha diversity indices (Supplementary figure 14). In contrast, no diarrhea-associated differences in gut microbiota age or microbial diversity was observed in infants who received more than two months of EBF (all p-values for heterogeneity tests <0.05).

Diarrhea at the time of sample collection was also associated with major perturbation in the gut bacterial composition of infants who received less than 2 months of EBF with a significant increase in the relative abundance of family Streptococcaceae and a significant decrease in the relative abundances of Bifidobacteriaceae and Coriobacteriaceae (all p-values <0.05 and FDR adjusted p-values <0.1). These changes in microbial composition were not observed in infants who received more than 2 months of EBF (Figure 5e, Supplementary table 12). Diarrhea at the time of sample collection was associated with an even more striking perturbation in the gut bacterial composition in infants who were not concurrently being breastfed, with a large outgrowth of Streptococcaceae and a tremendous decrease in the relative abundance of Bifidobactericeae (all p-values <0.05). These perturbations were almost absent in infants receiving breast milk at the time of diarrhea (Figure 5f, Supplementary table 12). The incidence of diarrhea was not different between breastfeeding statuses.

## Discussion

Our study analyzed data from seven microbiome studies and performed meta-analysis pooling estimates across studies with a total of 1825 stool samples of 684 infants from five countries. We found remarkably consistent differences between non-EBF vs. EBF infants in gut microbial diversity, microbiota age, microbial composition and microbial predicted functional pathways. The infants’ mode of delivery modified these effects. We also found notable interaction effects between breastfeeding and diarrhea on infant gut microbiota. With large datasets combined from different populations, our results are more robust and generalizable than from a single study.

Prior studies have reported increased bacterial species richness or diversity in non-EBF vs. EBF^28^ and/or trends of increased bacterial diversity across EBF, non-EBF and non-BF. ^29,31,32^ However, studies report different indices, analyze data in different ways and some do not account for age of infants at the time of stool sample collection, which is associated with breastfeeding status and infant gut microbiota. Our results showed a significant and consistent increase in all four commonly-used alpha diversity indices in non-EBF vs. EBF infants ≤ 6 months of age after adjusting for age of infants at sample collection. We also showed a consistent increase in gut microbiota age in non-EBF infants vs. EBF infants before 6 months of age across studies. At first glance these results may seem paradoxical but we hypothesize that the increase in diversity and microbiota age associated with non-EBF in the first 6 months may be out of step with age-appropriate development. A more stable, less diverse gut microbiota, associated with EBF, may be necessary in the early months of development.

There was substantial heterogeneity across studies and populations in gut bacterial taxonomic composition and gut bacterial metabolic pathway composition differences between non-EBF vs. EBF infants in the first 6 months of life. For example, the decrease in relative abundance of Proteobacteria in non-EBF vs. EBF infants was observed in four studies in North America but the opposite was observed in other studies in Bangladesh, Haiti and South Africa. This heterogeneity may be due to dietary differences or differences in formula ingredients in the non-EBF group across different populations. In addition, the gut microbiota of EBF infants, which is largely seeded by their mothers’ breastmilk microbiota and HMOs ^3,5,9,12^, might also be influenced by the mother’s diet or other exposures, which might also be different across populations ^10^. Infant ethnicity has been reported to influence the infant gut microbiota ^36^. In addition, variation in the region of 16S RNA gene targeted between studies may also contribute to heterogeneity. Despite this expected variation, our results revealed some important consistencies across populations. Our results showed a consistent increase in the relative abundance of Bacteroidetes in non-EBF vs. EBF infants in all seven studies as well as an overall significant increase in relative abundance of Firmicutes. More specifically, our results showed a persistent increase in relative abundances of genera *Bacteroides*, *Eubacterium* and *Veillonella* in non-EBF vs. EBF infants. While *Bifidobacterium* is the most common bacterial genus in gastrointestinal tract of young infants, *Bacteroides* and *Eubacterium* are the most common bacterial genera in the gastrointestinal tract of adults ^37,38^. Though these genera may be part of normal gut bacterial community, the increase in abundance of gut *Bacteroides* has been shown to be associated with higher body mass index (BMI) in young children ^39^ and *Veillonella* can be associated with different types of infection ^40^. Our results also showed a consistent increase in relative abundances of major microbial predicted pathways related to carbohydrate metabolism as well as a consistent decrease in relative abundances of crucial pathways related to lipid metabolism/homeostasis, free radical detoxification and metabolism of cofactors and vitamins in non-EBF infants ^41^. These findings may provide insight into biological mechanisms for the higher risk of obesity, diabetes and other adverse health outcomes in children who were not breastfed or non-exclusively breastfed in early months of life ^25–27^.

Interestingly, our results revealed notable heterogeneity regarding the perturbation in microbial-predicted functional pathways associated with non-EBF stratified by mode of delivery. The remarkably larger number of perturbed pathways of different cellular and metabolic processes in infants of cesarean deliveries may suggest that gut microbiota in these infants are more vulnerable to the effects of non-EBF. We also observed that non-EBF infants had a much lower abundance of Proteobacterial species than EBF infants after cesarean delivery. It appears that when gut microbiota are depleted with Bacteroidetes, as characteristically in early infancy after cesarean delivery, formula-feeding further depletes Proteobacteria ^42^. These findings may shed light on the mechanisms for the higher risk of adverse health outcomes in infants delivered by cesarean section ^43–45^ and emphasize the importance of EBF in cesarean-delivered infants. Differences in breastfeeding practices by mode of delivery may also account for these findings ^46^.

Effects of EBF in the first 6 months of life were still evident on the infant gut microbiota between 6 months to 2 years of age. Shorter duration of EBF was associated with increased gut microbiota age as well as earlier and larger increases in relative abundances of many bacterial families other than the beneficial family Bifidobacteriaceae. In contrast, longer EBF maintained a more stable bacterial composition in the early months of life and was associated with higher relative abundance of Bifidobacteriaceae. These findings again support our hypothesis that early changes in gut microbiota associated with non-EBF may be disproportional to immunological and biological maturity of infants in early months. That is, EBF nourishes a stable gut bacterial taxa composition that may be beneficial for the infants who are still immature in early month of life. Our results are also consistent with published literature that exposures in early life can affect the establishment of the gut microbiota in older children and adults ^36,47–49^ and may help explain the mechanism for the short and long-term health effects of EBF in early months of life.

Another particularly intriguing finding from our analysis is the protective effect of EBF on the infant gut microbiota during diarrheal episodes. Diarrhea has been previously shown to cause perturbations in the gut microbiota ^35,50^. Our analysis revealed that diarrhea was associated with a loss of microbial diversity, microbiota age and the relative abundance of Bifidobacteriaceae as well as an increase in the relative abundance of Streptococcaceae. Incredibly, these changes were almost completely abolished in infants who received more than 2 months of EBF as well as in those who were being breastfed at the time of diarrhea. These findings highlight the importance of longer duration of EBF in the first 6 months of life and continuation of breastfeeding after 6 months of life in maintaining a homeostatic gut microbiota that may be more resistant to outgrowth of pathogenic microbes that lead to diarrhea. Taken together, our findings suggest a key role for gut commensal bacteria in mediating the protective effects of breastfeeding on diarrhea morbidity and mortality.

In terms of methodology, our study applied appropriate and robust methodological approaches for the analysis of microbiome data and meta-analysis across microbiome studies. Standardization of alpha diversity indices and predicted microbiota age from each study makes the estimates of these measures comparable between studies. Zero-inflated beta GAMLSS models allow proper examination of relative abundances of bacterial taxa and predicted functional pathways, which range from zero to one and are generally zero-inflated as well as adjustment for confounding covariates and handling longitudinal or cross-sectional data. The estimates from zero-inflated beta GAMLSS models are log(odds ratio) of relative abundances and thus are comparable between studies. All effect estimates in our analyses were adjusted for variation in age of the infant at sample collection which might largely influence the infant gut microbiota composition as well as breastfeeding status but was not routinely accounted for in analysis ^3,29^ or was partially accounted by study design (collecting samples at similar infant age) 31,33 in some published studies. There have been some published meta-analyses for microbiome data ^51–56^ but none of these addressed between-group-comparison pooled effects when combining data from many studies as done here. The use of random effect meta-analysis models pooling estimates from studies allows examination of study-specific effects, the heterogeneity between studies, and the overall pooled effects across studies.

This study has some limitations. First, definitions of EBF and non-EBF were not identical across seven included studies. Specifically, for the five studies in Bangladesh, Canada, CA-FL, North Carolina and CA-MA-MO, EBF was defined as ingestion of only breastmilk without formula or solid food and non-EBF as ingestion of breastmilk plus formula and/or solid food. Whereas, the two studies in Haiti and South Africa followed World Health Organization (WHO) guidelines with EBF as feeding only breastmilk and non-EBF as feeding breast milk plus anything other than breast milk including traditional medicines or water. This difference might contribute to some of the variation in results across studies. Second, several of the studies were of a relatively small size or included other potential issues. Specifically, the Haiti study included only 48 infants, of which half were born to HIV-infected mothers, which has been shown to influence the infant gut microbiota ^3^. The North Carolina study included a very small number of samples in infants ≤ 6 months old (n=21). The VDAART trial (CA-MA-MO) study included samples from infants who were at high risk for asthma and allergies half of whom were randomized to vitamin D supplementation in pregnancy. However, sensitivity meta-analyses excluding these studies showed similar findings to the overall meta-analysis, suggesting that our results are robust. Finally, all of the results pertaining to samples collected after 6 months of age were from a single study (Bangladesh) and thus should be replicated in other cohorts. However, this study included nearly 1,000 samples collected monthly from birth to 2 years of life in 50 subjects with detailed meta-data ^35^.

## Conclusion

Our meta-analysis revealed robust findings across populations that may help elucidate the effects of EBF on the infant gut microbiota. Non-EBF or shorter duration of EBF in the first six months of life was associated with higher gut microbial diversity, higher microbiota age, bacterial composition more closely resembling the adult microbiota, higher composition of bacterial functional pathways related to carbohydrate metabolism and lower composition of bacterial functional pathways related to lipid metabolism, detoxification, cofactor and vitamin metabolism. The perturbation in microbial functional pathways associated with non-EBF was larger in cesarean-delivered infants than in vaginally-delivered infants. Furthermore, EBF, especially longer than two months from birth, was associated with a more stable gut bacterial taxa composition and reduced diarrhea-associated microbial dysbiosis. The early and large change associated with non-EBF in the infant gut microbiota may be disproportional to age-appropriate immunological and biological development of the infant. Altogether, our results highlight a consistent role of EBF to maintain a homeostatic developmental trajectory of the infant gut microbiota and shed light on the mechanisms of the short and long-term benefits of EBF in the first six months of life.

## Methods

### Data sources and study population

Processed and partially processed 16S rRNA gene sequence data of stool samples were obtained from seven previously published studies ^3,5,18,28,29,31,35^. Of the included studies, three were from the US, one from Canada, one from Haiti, one from South Africa and one from Bangladesh. There were five studies with three breastfeeding categories (EBF, non-EBF and non-BF) and two studies with two breastfeeding categories (EBF and non-EBF). The total number of samples of infants ≤ 6 months of age included in the overall meta-analyses was 1151 (EBF=547, non-EBF=436, non-BF=168). There were four studies with available information regarding the infants’ mode of delivery included in meta-analyses stratified by mode of delivery with a total number of samples of 670 (vaginal deliveries =484, cesarean deliveries =186). The Bangladesh study contained 674 samples from 6 months to 2 years of age. In total, 1825 samples of 684 infants were used in our analyses. A summary of the included studies, data characteristics and prior data processing are presented in Table 1.

### Data processing

Sequence data from each included study were processed separately. To achieve necessary data consistency for meta-analyses in this study, OTU picking was performed at 97% similarity using QIIME version 1.9.1 ^57^ with the Greengenes database (version 13.8) ^58^. Alpha rarefaction was done in QIIME using default options. Rarefaction depth was selected as the highest depth that retained all study samples. Taxonomic relative abundances from phylum to genus levels and alpha diversity indices were calculated based on rarefied OTU tables. Metagenomic functional compositions of stool bacterial communities were predicted based on the normalized OTU tables using PICRUSt ^59^ and relative abundances were then calculated for the resulting KEGG (Kyoto Encyclopedia of Genes and Genomes) functional pathways ^41^.

### Statistical analysis

For each study, mean alpha diversity indices were calculated for each sample at the selected rarefaction depth. Random Forest (RF) modeling of gut microbiota maturity has been widely used to characterize development of the microbiota over chronological time ^5,35,49^. Adapting the approach from Subramanian et al. ^35^, relative abundances of 36 bacterial genera (Supplementary table 1) that were detected in the data of all seven included studies were regressed against infant chronological age using a RF model on the training dataset of the Bangladesh study. The RF model fit based on relative abundances of these shared bacterial genera was then used to predict infant age on the test data of the Bangladesh study and the data of each other included study. The predicted infant age based on relative abundances of these shared bacterial genera in each study is referred to as gut microbiota age in this paper. Alpha diversity indices and microbiota age from each study were standardized to have a mean of 0 and standard deviation of 1 to make these measures comparable across studies. Generalized additive mixed model (GAMM) as well as linear mixed effect model with subject random intercept (for longitudinal data) or linear model (for non-longitudinal data) adjusted for infant age at the time of stool sample collection were used to further examine the curves of standardized alpha diversity indices and standardized gut microbiota age over age of infants as well as the difference between groups in each study.

For each study, the summary tables of bacterial taxa and pathway relative abundances were filtered to retain only the taxa and pathways that had an average relative abundance of at least 0.005% and were detected in at least 5% of the number of samples in that study. Relative abundances of bacterial taxa and bacterial KEGG metabolic pathways were examined using Generalized Additive Models for Location Scale and Shape (GAMLSS) with zero inflated beta family (BEZI) and (mu) logit links and other default options as implemented in the R package gamlss ^60^. This approach allows proper examination of microbiome relative abundance data, which range from zero to one and are generally zero-inflated, as well as adjustment for covariates (e.g. infant age at sample collection) and handling of longitudinal data by including a subject random effect. In each study, to roughly test for trends across three breastfeeding categories (EBF, non-EBF and non-BF), breastfeeding was coded as a continuous variable in the models.

To examine overall effects while addressing heterogeneity across studies, random effect meta-analysis models with inverse variance weighting and DerSimonian–Laird estimator for between-study variance were used to pool the adjusted estimates and their standard errors from all included studies. Meta-analyses were done for only bacterial taxa and pathways whose adjusted estimates and standard errors were available in at least 50% of the number of included studies. All statistical tests were two sided. P-values <0.05 were regarded as significant and false discovery rate (FDR) adjusted p-values <0.1 were regarded as significant after adjusting for multiple testing.

## Code availability

The R code used to generate the results in this paper are available at https://github.com/nhanhocu/metamicrobiome_breastfeeding.

## Data availability

All data used in the analyses of this study are included in these published articles and their supplementary information files (references ^3,5,18,28,29,31,35^). The data from the Bangladesh study were downloaded from: https://gordonlab.wustl.edu/Subramanian_6_14/Nature_2014_Processed_16S_rRNA_datasets.html. The data from six other studies were obtained directly from the investigators. All processed microbiome datasets and relevant meta-data from all included studies as well as intermediate saved outputs used to generate the results in this paper are available at https://github.com/nhanhocu/metamicrobiome_breastfeeding.

## Competing interests

The authors declare that they have no competing interests.

## Funding

This work was supported by Mervyn W. Susser fellowship in the Gertrude H. Sergievsky Center, Columbia University Medical Center (to N.T.H); NIH grant numbers U01HL091528 and R01HL108818 (to S.T.W and A.A.L); NICHD grant number K08HD069201, a developmental grant from the University of Washington Center for AIDS Research (CFAR), an NIH funded program under award number AI027757, and the following NIH Institutes and Centers: NIAID, NCI, NIMH, NIDA, NICHD, NHLBI, NIA, NIGMS, NIDDK (to H.B.J); NIH/NIDDK grant number P30 DK34987 (to M.A.A.P); NIH Grant Numbers K23 HD072774-02 (to P.S.P), K12 HD 052954-09 (to J.M.B), and UM1AI106716 (to G.M.A); grant 227312 from the Canadian Institutes of Health Research Canadian Microbiome Initiative (to A.L.K), the Canadian Institutes of Health Research and the Allergy, Genes, and Environment (AllerGen) Network of Centres of Excellence for the CHILD Study.

## Authors’ contribution

N.T.H and L.K conceived the study. F.L, K.A.L, H.M.T, B.B, P.S.P, L.F.W, J.M.B, J.E.S, M.B.A, A.L.T, S.T.W, M.A.A.P, A.A.L, A.L.K, H.B.J, G.M.A provided data. N.T.H performed the data analysis with inputs from F.L and L.K. N.T.H, F.L, H.B.J, G.M.A and L.K prepared the manuscript with inputs from all authors. All authors read and approved the final manuscript.

### Acknowledgement

We would like to acknowledge **CHILD Study investigators** for providing data of the study in Canada including**: Subbarao P** (Director), The Hospital for Sick Children & University of Toronto; **Turvey SE** (co-Director), University of British Columbia; **Sears MR**, (Founding Director), McMaster University; **Anand SS**, McMaster University; **Becker AB**, University of Manitoba; **Befus AD**, University of Alberta; **Brauer M**, University of British Columbia; **Brook JR**, University of Toronto; **Chen E**, Northwestern University, Chicago; **Cyr MM**, McMaster University; **Daley D**, University of British Columbia; **Dell SD**, The Hospital for Sick Children & University of Toronto; **Denburg JA**, McMaster University; **Duan QL**, Queen’s University; **Eiwegger T**, The Hospital for Sick Children & University of Toronto; **Grasemann H**, The Hospital for Sick Children & University of Toronto; **HayGlass K**, University of Manitoba; **Hegele RG**, The Hospital for Sick Children & University of Toronto; **Holness DL**, University of Toronto; **Hystad P**, Oregon State University; **Kobor M**, University of British Columbia; **Kollmann TR**, University of British Columbia; **Kozyrskyj AL**, University of Alberta; **Laprise C**, Université du Québec à Chicoutimi; **Lou WYW**, University of Toronto; **Macri J**, McMaster University; **Mandhane PJ**, University of Alberta; **Miller G**, Northwestern University, Chicago; **Moraes TJ**, The Hospital for Sick Children & University of Toronto; **Paré P**, University of British Columbia; **Ramsey C**, University of Manitoba; **Ratjen F**, The Hospital for Sick Children & University of Toronto; **Sandford A**, University of British Columbia; **Scott JA**, University of Toronto; **Scott J**, University of Toronto; **Silverman F**, University of Toronto; **Simons E**, University of Manitoba; **Takaro T**, Simon Fraser University; **Tebbutt SJ**, University of British Columbia; **To T**, The Hospital for Sick Children & University of Toronto.

## References

1. Rautava, S. Early microbial contact, the breast milk microbiome and child health. J. Dev. Orig. Health. Dis. 7, 5–14 (2016).

2. Nagata, R. et al. Transmission of the major skin microbiota, Malassezia, from mother to neonate. Pediatr. Int. 54, 350–355 (2012).

3. Bender, J. M. et al. Maternal HIV infection influences the microbiome of HIV-uninfected infants. Sci. Transl. Med. 8, 349ra100 (2016).

4. Schanche, M. et al. High-Resolution Analyses of Overlap in the Microbiota Between Mothers and Their Children. Curr. Microbiol. 71, 283–290 (2015).

5. Pannaraj, P. S. et al. Association Between Breast Milk Bacterial Communities and Establishment and Development of the Infant Gut Microbiome. JAMA Pediatr. 90095, 647–654 (2017).

6. Newburg, D. S. & Morelli, L. Human milk and infant intestinal mucosal glycans guide succession of the neonatal intestinal microbiota. Pediatr. Res. 77, 115–20 (2015).

7. Kozak, K., Charbonneau, D., Sanozky-Dawes, R. & Klaenhammer, T. Characterization of bacterial isolates from the microbiota of mothers’ breast milk and their infants. Gut Microbes 6, 341–351 (2015).

8. Wang, M. et al. Fecal Microbiota Composition of Breast-Fed Infants Is Correlated With Human Milk Oligosaccharides Consumed. J. Pediatr. Gastroenterol. Nutr. 60, 825–833 (2015).

9. Bashiardes, S., Thaiss, C. A. & Elinav, E. It’s in the milk: Feeding the microbiome to promote infant growth. Cell Metab. 23, 393–394 (2016).

10. Cabrera-Rubio, R. et al. The human milk microbiome changes over lactation and is shaped by maternal weight and mode of delivery. Am. J. Clin. Nutr. 96, 544–551 (2012).

11. González, R. et al. Breast milk and gut microbiota in African mothers and infants from an area of high HIV prevalence. PLoS One 8, e80299 (2013).

12. Davis, J. C. C. et al. Identification of Oligosaccharides in Feces of Breast-fed Infants and Their Correlation with the Gut Microbial Community. Mol. Cell. Proteomics 15, 2987–3002 (2016).

13. Milani, C. et al. The First Microbial Colonizers of the Human Gut: Composition, Activities, and Health Implications of the Infant Gut Microbiota. Microbiol. Mol. Biol. Rev. 81, e00036–17 (2017).

14. Azad, M. B. et al. Gut microbiota of healthy Canadian infants: profiles by mode of delivery and infant diet at 4 months. CMAJ 185, 385–94 (2013).

15. Fan, W. et al. Diversity of the intestinal microbiota in different patterns of feeding infants by Illumina high-throughput sequencing. World J. Microbiol. Biotechnol. 29, 2365–2372 (2013).

16. Gomez-Llorente, C. et al. Three main factors define changes in fecal microbiota associated with feeding modality in infants. J. Pediatr. Gastroenterol. Nutr. 57, 461–6 (2013).

17. Gregory, K. E. et al. Influence of maternal breast milk ingestion on acquisition of the intestinal microbiome in preterm infants. Microbiome 4, 68 (2016).

18. Sordillo, J. E. et al. Factors influencing the infant gut microbiome at age 3-6 months: Findings from the ethnically diverse Vitamin D Antenatal Asthma Reduction Trial (VDAART). J. Allergy Clin. Immunol. 139, 482–491.e14 (2017).

19. Timmerman, H. M. et al. Intestinal colonisation patterns in breastfed and formula-fed infants during the first 12 weeks of life reveal sequential microbiota signatures. Sci. Rep. 7, 8327 (2017).

20. Tsiotsias, A. & Welling, G. W. Microbiota profile in feces of breast- and formula-fed newborns by using fluorescence in situ hybridization (FISH). Anaerobe 17, 478–482 (2011).

21. Davis, M. Y., Zhang, H., Brannan, L. E., Carman, R. J. & Boone, J. H. Rapid change of fecal microbiome and disappearance of Clostridium difficile in a colonized infant after transition from breast milk to cow milk. Microbiome 4, 53 (2016).

22. Kramer, M. S. & Kakuma, R. in Cochrane Database of Systematic Reviews (ed. Kramer, M. S.) (John Wiley & Sons, Ltd, 2012). doi:10.1002/14651858.CD003517.pub2

23. WHO | Exclusive breastfeeding for six months best for babies everywhere. WHO (2011).

24. Lamberti, L. M., Fischer Walker, C. L., Noiman, A., Victora, C. & Black, R. E. Breastfeeding and the risk for diarrhea morbidity and mortality. BMC Public Health 11, S15 (2011).

25. Stuebe, A. The risks of not breastfeeding for mothers and infants. Rev. Obstet. Gynecol. 2, 222–31 (2009).

26. Yan, J., Liu, L., Zhu, Y., Huang, G. & Wang, P. P. The association between breastfeeding and childhood obesity: a meta-analysis. BMC Public Health 14, 1267 (2014).

27. Cardwell, C. R. et al. Breast-feeding and childhood-onset type 1 diabetes: a pooled analysis of individual participant data from 43 observational studies. Diabetes Care 35, 2215–25 (2012).

28. Wood, L. et al. Feeding mode regulates gut microbial composition, peripheral T cell activation and mucosal gene expression in African infants. Clin. Infect. Dis. In press, (2018).

29. Thompson, A. L., Monteagudo-Mera, A., Cadenas, M. B., Lampl, M. L. & Azcarate-Peril, M. A. Milk- and solid-feeding practices and daycare attendance are associated with differences in bacterial diversity, predominant communities, and metabolic and immune function of the infant gut microbiome. Front. Cell. Infect. Microbiol. 5, 3 (2015).

30. Bokulich, N. A. et al. Antibiotics, birth mode, and diet shape microbiome maturation during early life. Sci. Transl. Med. 8, 343ra82 (2016).

31. Azad, M. B. et al. Impact of maternal intrapartum antibiotics, method of birth and breastfeeding on gut microbiota during the first year of life: A prospective cohort study. BJOG: An International Journal of Obstetrics and Gynaecology 123, 983–993 (2016).

32. Hesla, H. M. et al. Impact of lifestyle on the gut microbiota of healthy infants and their mothers—the ALADDIN birth cohort. FEMS Microbiol. Ecol. 90, 791–801 (2014).

33. Madan, J. C. et al. Association of Cesarean Delivery and Formula Supplementation With the Intestinal Microbiome of 6-Week-Old Infants. JAMA Pediatr. 170, 212 (2016).

34. Chu, D. M. et al. Maturation of the infant microbiome community structure and function across multiple body sites and in relation to mode of delivery. Nat. Med. 23, 314–326 (2017).

35. Subramanian, S. et al. Persistent gut microbiota immaturity in malnourished Bangladeshi children. Nature 510, 417–421 (2014).

36. Stearns, J. C. et al. Ethnic and diet-related differences in the healthy infant microbiome. Genome Med. 9, 32 (2017).

37. Lagier, J.-C., Million, M., Hugon, P., Armougom, F. & Raoult, D. Human gut microbiota: repertoire and variations. Front. Cell. Infect. Microbiol. 2, 136 (2012).

38. Schwiertz, A., Le Blay, G. & Blaut, M. Quantification of different Eubacterium spp. in human fecal samples with species-specific 16S rRNA-targeted oligonucleotide probes. Appl. Environ. Microbiol. 66, 375–82 (2000).

39. Vael, C., Verhulst, S. L., Nelen, V., Goossens, H. & Desager, K. N. Intestinal microflora and body mass index during the first three years of life: an observational study. Gut Pathog. 3, 8 (2011).

40. Brook, I. Veillonella Infections in Children. J. Clin. Microbiol. 34, 1283–1285 (1996).

41. KEGG PATHWAY Database. Available at: http://www.genome.jp/kegg/pathway.html. (Accessed: 22nd November 2017)

42. Chua, M. C. et al. Effect of Synbiotic on the Gut Microbiota of Cesarean Delivered Infants: A Randomized, Double-blind, Multicenter Study. J. Pediatr. Gastroenterol. Nutr. 65, 102–106 (2017).

43. Black, M., Bhattacharya, S., Philip, S., Norman, J. E. & McLernon, D. J. Planned Cesarean Delivery at Term and Adverse Outcomes in Childhood Health. JAMA 314, 2271 (2015).

44. Neu, J. & Rushing, J. Cesarean versus Vaginal Delivery: Long term infant outcomes and the Hygiene Hypothesis. doi:10.1016/j.clp.2011.03.008

45. Yasmin, F. et al. Cesarean Section, Formula Feeding, and Infant Antibiotic Exposure: Separate and Combined Impacts on Gut Microbial Changes in Later Infancy. Front. Pediatr. 5, 1–13 (2017).

46. Hobbs, A. J., Mannion, C. A., Mcdonald, S. W., Brockway, M. & Tough, S. C. The impact of caesarean section on breastfeeding initiation, duration and difficulties in the first four months postpartum. BMC Pregnancy Childbirth 16, (2016).

47. Lemas, D. J. et al. Exploring the contribution of maternal antibiotics and breastfeeding to development of the infant microbiome and pediatric obesity. Semin. Fetal Neonatal Med. 21, 406–409 (2016).

48. Laursen, M. F., Bahl, M. I., Michaelsen, K. F. & Licht, T. R. First Foods and Gut Microbes. Front. Microbiol. 8, 356 (2017).

49. Bäckhed, F. et al. Dynamics and stabilization of the human gut microbiome during the first year of life. Cell Host Microbe 17, 690–703 (2015).

50. The, H. C. et al. Assessing gut microbiota perturbations during the early phase of infectious diarrhea in Vietnamese children. Gut Microbes 00–00 (2017). doi:10.1080/19490976.2017.1361093

51. Adams, R. I., Bateman, A. C., Bik, H. M. & Meadow, J. F. Microbiota of the indoor environment: a meta-analysis. 3, (2015).

52. Bhute, S. et al. Molecular Characterization and Meta-Analysis of Gut Microbial Communities Illustrate Enrichment of Prevotella and Megasphaera in Indian Subjects. Front. Microbiol. 7, 660 (2016).

53. Holman, D. B., Brunelle, B. W., Trachsel, J. & Allen, H. K. Meta-analysis To Define a Core Microbiota in the Swine Gut. mSystems 2, (2017).

54. Mancabelli, L. et al. Meta-analysis of the human gut microbiome from urbanized and pre-agricultural populations. Environ. Microbiol. 19, 1379–1390 (2017).

55. Lozupone, C. A. et al. Meta-analyses of studies of the human microbiota. Genome Res.23, 1704–14 (2013).

56. Duvallet, C., Gibbons, S. M., Gurry, T., Irizarry, R. A. & Alm, E. J. Meta-analysis of gut microbiome studies identifies disease-specific and shared responses. Nat. Commun. 8, 1784 (2017).

57. Caporaso, J. G. et al. QIIME allows analysis of high-throughput community sequencing data. Nat. Methods 7, 335–336 (2010).

58. DeSantis, T. Z. et al. Greengenes, a chimera-checked 16S rRNA gene database and workbench compatible with ARB. Appl. Environ. Microbiol. 72, 5069–72 (2006).

59. Langille, M. G. I. et al. Predictive functional profiling of microbial communities using 16S rRNA marker gene sequences. Nat. Biotechnol. 31, 814–821 (2013).

60. Rigby, R. A. & Stasinopoulos, D. M. Generalized additive models for location, scale and shape (with discussion). J. R. Stat. Soc. Ser. C (Applied Stat. 54, 507–554 (2005).

